# MotilA – A Python pipeline for the analysis of microglial fine process motility in 3D time-lapse multiphoton microscopy data

**DOI:** 10.1101/2025.08.04.668426

**Authors:** Fabrizio Musacchio, Sophie Crux, Felix Nebeling, Nala Gockel, Falko Fuhrmann, Martin Fuhrmann

**Affiliations:** Neuroimmunology & Imaging group, German Center for Neurodegenerative Diseases (DZNE), Bonn, Germany

**Author notes:** Corresponding authors: Dr. Fabrizio Musacchio. Prof. Dr. Martin Fuhrmann, Neuroimmunology and Imaging Group, German Center for Neurodegenerative Diseases (DZNE), Venusberg-Campus 1/99, 53127 Bonn, Germany.

## Abstract

*MotilA* is a Python-based image analysis pipeline for quantifying fine process motility of microglia from 3D time-lapse two-channel fluorescence microscopy data. Developed for high-resolution multiphoton *in vivo* imaging datasets, *MotilA* enables both single-file and batch processing across multiple experimental conditions. It performs image preprocessing, segmentation, and motility quantification over time, using a pixel-based change detection strategy that yields biologically interpretable metrics such as the turnover rate (TOR) of microglial fine processes. While originally designed for microglial imaging, the pipeline can be extended to other cell types and imaging applications that require analysis of dynamic morphological changes. *MotilA* is openly available, platform-independent, and includes extensive documentation, tutorials, and example data to facilitate adoption by the broader scientific community. It is released under the GPL-3.0 open-source license.

## Statement of need

Microglia are innate immune cells of the central nervous system and exhibit highly dynamic, motile processes that continuously scan their environment (Nimmerjahn, Kirchhoff, and Helmchen 2005; M. Fuhrmann et al. 2010; Tremblay, Lowery, and Majewska 2010; Prinz, Jung, and Priller 2019; Nebeling et al. 2023). Quantifying microglial motility at the level of fine processes is crucial for studying their function in health and disease, including neurodegeneration, inflammation, and synaptic remodeling. However, despite the biological importance of this analysis, there is currently no dedicated open-source tool tailored for this task.

To date, researchers typically quantify microglial motility manually (see., e.g., Nebeling et al. (2023)) using general-purpose image processing software such as Fiji/ImageJ (Schindelin et al. 2012) or ZEISS ZEN (Carl Zeiss Microscopy GmbH Accessed 2025). While these approaches are well established in the field, they are time-consuming, lack reproducibility, and are not well suited for batch processing or cohort-level comparisons. They often focus on individual microglia, whereas *MotilA* enables analysis of the full field of view, allowing for more comprehensive and scalable quantification. Manual workflows are also more susceptible to human bias, limiting their scalability and objectivity (Lee et al. 2024; Wall et al. 2018; Misra et al. 2015; Brown 2017).

*MotilA* addresses these limitations by providing an end-to-end, user-friendly, and batch-capable pipeline specifically designed for 3D time-lapse two-channel microscopy data. It supports standardized workflows for single- and multi-channel datasets, integrates essential preprocessing steps (registration, spectral unmixing, histogram normalization), and derives biologically meaningful motility metrics from binarized pixel dynamics. The method builds on strategies used in prior studies but automates the workflow in a reproducible, scalable, and open-source manner. *MotilA* thus fills a critical gap in neuroimaging analysis pipelines and is particularly valuable for labs working with multiphoton *in vivo* imaging.

### What does *MotilA* do?

*MotilA* is a modular and customizable image analysis pipeline written in Python that quantifies microglial fine process motility from time-lapse fluorescence microscopy data, typically acquired with two-photon (Denk, Strickler, and Webb 1990; Helmchen and Denk 2005) or three-photon *in vivo* imaging (Horton et al. 2013; F. Fuhrmann et al. 2024). Although it was originally developed for microglial analysis, the pipeline is adaptable to other cell types and imaging contexts involving dynamic morphological changes over time.

At its core, *MotilA* extracts sub-volumes from 3D time-stacks, performs 2D maximum intensity projections, and segments the resulting images to classify pixel-wise changes in microglial morphology. These changes are quantified frame-by-frame and used to calculate biologically interpretable metrics, including the turnover rate (TOR). The design is tailored to biological imaging data, with particular attention to typical issues such as z-axis projection loss, channel bleed-through, motion artifacts, photobleaching, and signal heterogeneity.

The pipeline supports both single-file processing and large-scale batch analysis. Parameters are highly customizable either programmatically or via metadata files, and all results are automatically logged, saved, and summarized for downstream statistical analysis. The outputs include segmented image series, intermediate diagnostics (e.g. histograms, projections, brightness traces), and Excel spreadsheets with motility metrics.

To accommodate large-scale, high-resolution imaging data, *MotilA* supports memory-efficient file handling via the Zarr format (Miles et al. 2025), enabling processing of large TIFF files using memory mapping to avoid RAM overload.

*MotilA* can be run via Python scripts or Jupyter notebooks, and it includes extensive documentation, examples, and a tutorial dataset to make onboarding straightforward.

We welcome community contributions and issue reports via the GitHub repository: https://github.com/FabrizioMusacchio/motila.

### How is “motility” determined?

*MotilA* quantifies motility by analyzing pixel-wise changes in microglial fine processes’ morphology over time. The pipeline first extracts a sub-volume around a user-defined z-axis center from each 3D image stack and applies a 2D maximum intensity projection to reduce dimensionality. Although this sacrifices some z-axis information, it enables efficient segmentation and pixel-level tracking while maintaining biological interpretability.

At each time point *t*_*i*_, the projected and binarized image *B*(*t*_*i*_) is compared to the next time point *B*(*t*_*i*+1_). A temporal variation map *ΔB*(*t*_*i*_) is computed as:

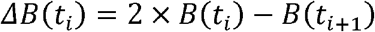

Based on this difference image, each pixel is classified as:

- **Stable (S)** if *ΔB* = 1
- **Gained (G)** if *ΔB* = -1
- **Lost (L)** if *ΔB* = 2

From these categories, *MotilA* calculates the microglial **fine process turnover rate (**TOR), a central metric representing the fraction of pixels that changed:

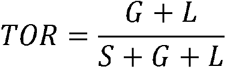

This approach allows for a quantitative assessment of microglial process dynamics at each time point and across the full recording session. The same principle can be extended to other motile cell types or dynamic cellular structures where morphological changes manifest as gain or loss of segmented pixels over time.

The implementation is based on analytical strategies described in previous studies such as M. Fuhrmann et al. (2010) and Nebeling et al. (2023), with added flexibility for batch processing, filtering, and parameter tuning.

### Key features

*MotilA* offers a combination of modularity, reproducibility, and scalability specifically tailored to motility analysis in multiphoton *in vivo* imaging. Its key features include:

- **Automated preprocessing pipeline** Includes optional steps for image registration (2D and 3D), spectral unmixing, histogram equalization for contrast enhancement within time points, and histogram matching for brightness normalization across time points (e.g. to correct for photobleaching), as well as noise reduction via median and Gaussian filtering.
- **Flexible segmentation and thresholding** Supports multiple adaptive thresholding methods (e.g. Otsu, Li, Triangle) and customizable blob detection settings to isolate fine microglial processes or similar structures.
- **Pixel-based motility quantification** Tracks pixel-wise changes between time points to classify stable, gained, and lost pixels, allowing biologically interpretable metrics like the turnover rate (TOR).
- **Batch processing capabilities** Enables large-scale processing of multiple datasets with a standardized folder structure and parameter metadata sheets, suitable for cohort-level studies.
- **User-defined projection settings** Allows flexible extraction of sub-volumes and z-projection around multiple centers to avoid overlapping cells and vascular artifacts.
- **Memory-efficient file handling** Supports memory mapping of large TIFF files via the Zarr format, enabling efficient processing of high-resolution time-lapse datasets without exhausting system RAM.
- **Metadata integration and parameter logging** Automatically reads per-dataset settings from metadata files (e.g. Excel sheets), and stores processing parameters and outputs in structured result folders.
- **Cross-platform compatibility** Runs on Windows, macOS, and Linux, tested with Python ≥3.9 and compatible with common scientific computing environments via Conda.
- **Tutorials and example data included**

Comes with Jupyter notebooks, example datasets, and clear documentation to help new users get started quickly.

### Pipeline steps

The *MotilA* pipeline follows a modular sequence of image processing and analysis steps designed for robust and reproducible quantification of motility from multi-dimensional imaging data. It supports both single-file and batch workflows and includes options for fine-grained customization at each step.

### Core pipeline steps

For single datasets, *MotilA* executes the following sequence (**Figure 1**)):

**Figure 1:**
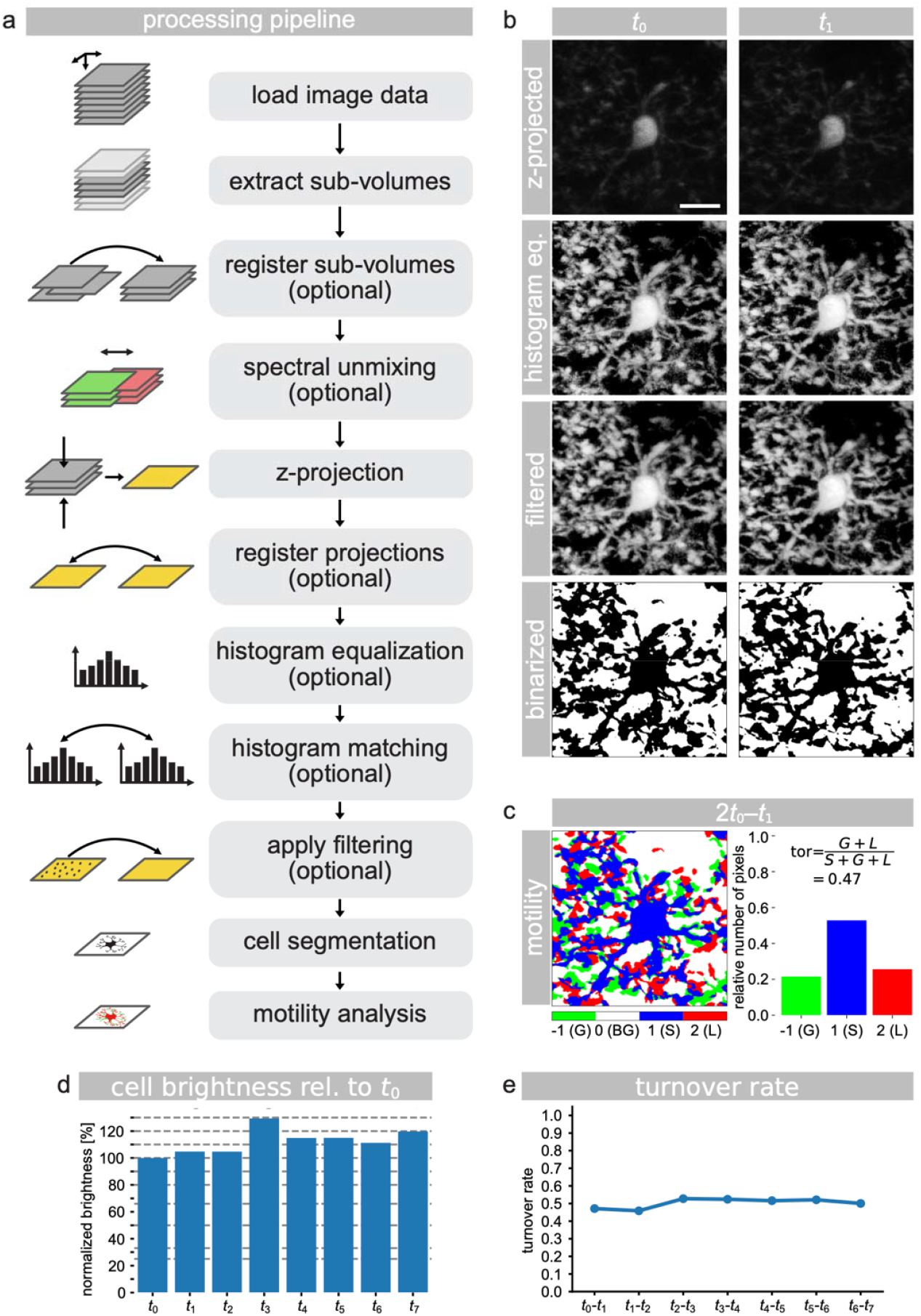
Step-by-step illustration of the MotilA pipeline using the included test dataset. **a)** Overview of the image processing pipeline, showing core and optional steps. **b)** Example projections of a cropped microglial cell at time points and, including raw, histogram-equalized, filtered (median and Gaussian), and binarized versions. **c)** Binarized pixel-wise comparison between and, with classification into stable (S, blue), gained (G, green), lost (L, red), and background (BG, white) pixels, along with the corresponding pixel statistics. **d)** Normalized cell brightness over time, relative to, used to assess bleaching and signal stability. **e)** Turnover rate (TOR) plotted across all time points for the same cell, representing process-level motility dynamics. All microglial image panels in b and c are shown at the same scale. Scale bar in the top-left image of panel b represents 10□μm.

1. **Load image data** Supports TIFF files (Gohlke 2025) in TZCYX (multi-channel) or TZYX (single-channel) format, following ImageJ/Fiji conventions (T: time, Z: z-axis, C: channel, Y: height, X: width).
2. **Extract sub-volumes** Selects a z-stack around a projection center for each time point, allowing focused analysis and optional multiple projections per stack.
3. **(Optional) Register sub-volumes** Applies 3D motion correction (Guizar-Sicairos, Thurman, and Fienup 2008; Anuta 1970; Kuglin and Hines 1975) to each time series stack using user-defined template strategies (e.g. mean, median).
4. **(Optional) Perform spectral unmixing** Removes signal bleed-through between channels, especially relevant for two-channel imaging setups.
5. **Z-projection** Projects each 3D sub-volume into a 2D image via maximum intensity projection to simplify segmentation and speed up processing.
6. **(Optional) Register projections** Aligns the 2D projections across time to correct for lateral motion artifacts.
7. **(Optional) Apply histogram equalization** Enhances local contrast within each projection using contrast-limited adaptive histogram equalization (CLAHE) (Pizer et al. 1987; Walt et al. 2014).
8. **(Optional) Apply histogram matching** Normalizes brightness across time points to mitigate bleaching effects or intensity drift (Walt et al. 2014).
9. **(Optional) Apply filtering** Reduces noise with optional median filtering (square or circular kernel) and/or Gaussian smoothing (Virtanen et al. 2020; Harris et al. 2020).
10. **Segment microglial processes** Applies adaptive thresholding (Ridler, Calvard, and name 1978; Otsu 1979; Li and Tam 1998; Glasbey 1993; Prewitt and Mendelsohn 1966; Zack, Rogers, and Latt 1977; Yen, Chang, and Chang 1995) and blob filtering (Fiorio and Gustedt 1996; Wu, Otoo, and Shoshani 2005; Walt et al. 2014) to identify and isolate morphologically relevant structures.
11. **Analyze motility** Quantifies pixel-level changes over time to classify stable, gained, and lost regions, from which motility metrics are derived.

All intermediate outputs and metrics are saved for validation and further analysis.

### Batch processing steps

*MotilA* supports fully automated batch processing using a standardized folder structure and Excel-based metadata configuration. This enables reproducible cohort-level analysis across many animals or experimental conditions.

1. **Define a project folder** Each dataset is placed in an ID-specific subdirectory, containing imaging files, metadata, and optional result directories.
2. **Run the batch process** The core pipeline is executed for each dataset using shared or per-dataset parameters defined in metadata.xls.
3. **Save results** Segmentation outputs, projections, and motility metrics are stored in structured result folders for each dataset.
4. **Batch-collect metrics**

Aggregates metrics across datasets into cohort-level Excel files for downstream statistical analysis.

This design enables large-scale, reproducible quantification of microglial motility with minimal manual intervention.

### Assessing results and analyzing outputs

*MotilA* provides rich output in the form of diagnostic plots, intermediate image files, and structured Excel tables to support both per-dataset assessment and cohort-level statistical analysis.

#### Per-dataset assessment

For each processed image stack, *MotilA* generates:

- **Segmented images and overlays** showing gained, lost, and stable regions across time points.
- **Histogram plots** for brightness, pixel area, and thresholding diagnostics.
- **Motility metrics table (motility.xlsx)** containing:
  ∘ Gained pixels (G)
  ∘ Lost pixels (L)
  ∘ Stable pixels (S)
  ∘ turnover rate (TOR) per time point
- **Brightness metrics** (brightness.xlsx) tracking average pixel intensity over time.
- **Cell area metrics** (cell_pixel_area.xlsx) reporting the segmented microglial pixel area per time point.

These outputs help assess segmentation quality, evaluate photobleaching or signal loss, and refine preprocessing parameters as needed.

#### Cohort-level batch analysis

During batch processing, *MotilA* can aggregate key metrics from all datasets into shared summary files, including:

- all_motility.xlsx — All G/L/S/TOR metrics across datasets.
- all_brightness.xlsx — Mean brightness per dataset and time point.
- all_cell_pixel_area.xlsx — Segmented area per dataset and time point.
- average_motility.xlsx — Dataset-wise average motility metrics across the full recording.

These results allow for statistical comparison of motility dynamics across experimental conditions and facilitate downstream visualization and modeling in tools like Python, R, or Excel.

This multi-level output strategy ensures both technical validation and biological insight, making *MotilA* suitable for both exploratory and hypothesis-driven studies.

### Main functions

The three main entry points for the pipeline are:

- process_stack — Processes a single image stack, performing the full pipeline.
- batch_process_stacks — Executes the pipeline across multiple datasets in a project folder.
- batch_collect — Gathers motility metrics from all datasets for cohort-level analysis. Each function supports extensive parameterization via arguments or metadata files.

A complete overview of configurable parameters for single-file processing, batch workflows, and image enhancement is provided in the MotilA README.

### Useful helper functions

Several additional functions assist with data preparation and quality control, including:

- tiff_axes_check_and_correct — Automatically adjusts TIFF axis order to TZCYX/ TZYX if needed.
- hello_world — Verifies a successful import of the *MotilA* module.
- logger_object — Initializes logging for the current analysis session.

### Applications and limitations

*MotilA* was designed with a primary focus on the analysis of microglial fine process motility *in vivo*, using high-resolution 3D time-lapse two-channel fluorescence microscopy data. Its modular design and general image processing framework, however, make it applicable to a broader range of dynamic imaging contexts.

### Applications

- **Microglial dynamics** Quantification of process turnover during surveillance, neuroinflammation, or disease models such as neurodegeneration and injury.
- **Neuronal structural plasticity** While *MotilA* is optimized for microglial processes, its pixel-based change detection framework can in principle be adapted to analyze dynamic changes in dendrites or axons — such as growth, retraction, or remodeling — provided the structures can be reliably segmented across time.
- **Two-channel *in vivo* imaging** Effective for experiments involving simultaneous imaging of microglia and neurons (e.g., Cx3Cr1-GFP with Thy1-YFP), with spectral unmixing to reduce bleed-through from overlapping channels or fluorophores.
- **Cohort-level studies** Designed to analyze and compare motility metrics across large experimental groups, enabling high-throughput, statistically robust results.
- **Teaching and prototyping** The example datasets and tutorials make *MotilA* a useful tool for training purposes or prototyping new analysis approaches.

### Limitations

- **Loss of z-resolution** The use of 2D maximum intensity projections simplifies processing but sacrifices z-axis information. This may lead to overlapping structures and limits spatial specificity.
- **Segmentation-dependent** Accuracy depends on appropriate thresholding and image quality. Overlapping processes, blood vessels, or low signal-to-noise ratios can reduce segmentation performance.
- **Limited spectral unmixing** The current unmixing approach is a simple channel subtraction. More advanced unmixing strategies may be required for some experimental setups.
- **Not a general-purpose tracking tool** *MotilA* is optimized for pixel-level process motility, not for full cell tracking or object-based morphological quantification over time.
- **Assumes TIFF input with standardized axis order.** Input images must conform to TZCYX or TZYX structure; other formats require conversion.

Despite these limitations, *MotilA* provides a powerful, reproducible framework for analyzing microglial motility and similar biological processes, especially in experimental setups where manual analysis would be impractical.

### Real-world example

To demonstrate its practical utility, *MotilA* includes a fully compatible example dataset of *in vivo* two-photon time-lapse imaging stacks from the mouse frontal cortex (Gockel et al. 2025). These data were acquired to assess microglial fine process motility under control conditions and during complement C4 overexpression, a genetic risk factor for schizophrenia.

The dataset contains two 5D TIFF stacks with the following structure:

- **T**: 8 time points (5-minute intervals over 35 minutes)
- **C**: 2 imaging channels (Cx3cr1-GFP for microglia, tdTomato for neurons)
- **Z**: ∼60 optical sections (1□μm step size)
- **Y, X**: ∼1200 × 1200 px (∼125□×□125□μm^2^ field of view)

The files are formatted for direct use with *MotilA*, requiring no manual reorganization or preprocessing.

Figure 1. summarizes the full *MotilA* workflow as applied to the example dataset. For visualization purposes, the original full-field dataset was cropped around a single microglial cell to reduce background clutter and allow detailed inspection of each processing step. **Panel a)** outlines the core and optional steps in the processing pipeline. **Panel b)** shows z-projections of the microglial cell at time points *t*_0_ and *t*_1_, including raw data, contrast enhancement, and filtering prior to segmentation. **Panel c)** displays the delta image used for motility quantification and the corresponding pixel-wise classification into stable (S, blue), gained (G, green), and lost (L, red) pixels. **Panel d)** tracks the average brightness of the segmented cell relative to the first time point, which helps assess signal stability and potential bleaching. **Panel e)** presents the turnover rate (TOR) across all time points, capturing the dynamics of microglial process remodeling.

### Past and ongoing projects

*MotilA* has already been successfully applied in multiple neuroscience studies involving *in vivo* imaging of microglia and neurons in the mouse brain.

The following published and preprint works used *MotilA* to analyze fine process motility in physiological and pathological contexts:

- **Crux et al. (2024)** Investigated the role of actin depolymerizing factors ADF/Cofilin1 in microglial motility and memory formation. *MotilA* was used to quantify reduced motility in knockout mice. → https://doi.org/10.1101/2024.09.27.615114
- **F. Fuhrmann et al. (2024)** Employed deep three-photon imaging of microglia in the medial prefrontal cortex to measure sub-cellular process dynamics in awake mice. *MotilA* was used to quantify microglial turnover at depths beyond 1 mm. → https://doi.org/10.1101/2024.08.28.610026
- **Gockel et al. (2025)** Generated and published the example dataset accompanying this pipeline, which was used to demonstrate microglial motility changes in response to complement C4 overexpression. → https://doi.org/10.5281/zenodo.15061566

These studies showcase the pipeline’s suitability for both targeted microglial investigations and large-scale, high-resolution imaging projects. Ongoing work continues to extend *MotilA*’s application to additional brain regions, genetic perturbations, and imaging modalities, including multi-channel and high-speed two-photon datasets.

## Acknowledgements

We gratefully acknowledge the **Light Microscopy Facility (LMF)** and **Animal Research Facility (ARF)** at the German Center for Neurodegenerative Diseases (DZNE), Bonn, for their essential support in data acquisition and technical infrastructure.

This work was supported by the DZNE and by grants to MF from the European Union ERC-CoG (MicroSynCom 865618) and the German Research Foundation DFG (SFB1089 C01, B06; SPP2395). MF is a member of the DFG Excellence Cluster ImmunoSensation2. This work was also supported by the iBehave network to MF and the CANTAR (CANcerTARgeting) network to FN, both funded by the Ministry of Culture and Science of the State of North Rhine-Westphalia. The funders had no role in study design, data collection and interpretation, or the decision to submit the work for publication. FN received additional funding from the Mildred-Scheel School of Oncology Cologne-Bonn.

All animal procedures related to the example dataset were conducted in compliance with institutional, national, and international regulations. Experiments were approved by the relevant animal care and use committees at DZNE (Germany), following guidelines equivalent to the ARRIVE 2.0 framework. All efforts were made to reduce the number of animals used and to refine experimental conditions in accordance with the 3Rs (Replacement, Reduction, and Refinement) principles.

We also acknowledge the open-source community whose tools and contributions made the development of *MotilA* possible.

## References

Anuta, Paul E. 1970. “Spatial Registration of Multispectral and Multitemporal Digital Imagery Using Fast Fourier Transform Techniques.” IEEE Transactions on Geoscience Electronics 8 (4): 353–68. 10.1109/TGE.1970.271435.

Brown, Danielle L. 2017. “Bias in Image Analysis and Its Solution: Unbiased Stereology.” Journal of Toxicologic Pathology 30 (3): 183–91. 10.1293/tox.2017-0013.

Carl Zeiss Microscopy GmbH. Accessed 2025. “ZEISS ZEN Microscopy Software.” https://www.zeiss.com/metrology/en/software/zeiss-zen-core.html.

Crux, Sophie, Marie Denise Roggan, Stefanie Poll, Felix C. Nebeling, Juliane Schiweck, Manuel Mittag, Fabrizio Musacchio, et al. 2024. “Deficiency of Actin Depolymerizing Factors ADF/Cfl1 in Microglia Decreases Motility and Impairs Memory.” bioRxiv. 10.1101/2024.09.27.615114.

Denk, Winfried, James H. Strickler, and Watt W. Webb. 1990. “Two-Photon Laser Scanning Fluorescence Microscopy.” Science 248 (4951): 73–76. 10.1126/science.2321027.

Fiorio, Christophe, and Jens Gustedt. 1996. “Two Linear Time Union-Find Strategies for Image Processing.” Theoretical Computer Science 154: 165–81. 10.1016/0304-3975(94)00262-2.

Fuhrmann, Falko, Felix C. Nebeling, Fabrizio Musacchio, Manuel Mittag, Stefanie Poll, Monika Müller, Eleonora Ambrad Giovannetti, et al. 2024. “Three-Photon in Vivo Imaging of Neurons and Glia in the Medial Prefrontal Cortex with Sub-Cellular Resolution.” bioRxiv. 10.1101/2024.08.28.610026.

Fuhrmann, Martin, Tobias Bittner, Christian K. E. Jung, Steffen Burgold, Richard M. Page, Gerda Mitteregger, Christian Haass, Frank M. LaFerla, Hans Kretzschmar, and Jochen Herms. 2010. “Microglial Cx3cr1 Knockout Prevents Neuron Loss in a Mouse Model of Alzheimer’s Disease.” Nature Neuroscience 13 (4): 411–13. 10.1038/nn.2511.

Glasbey, Chris A. 1993. “An Analysis of Histogram-Based Thresholding Algorithms.” CVGIP: Graphical Models and Image Processing 55 (6): 532–37. 10.1006/cgip.1993.1040.

Gockel, Nala, Nayadoleni Nieves-Rivera, Mélanie Druart, Külli Jaako, Falko Fuhrmann, Rebeka Rožkalne, Fabrizio Musacchio, et al. 2025. “Example Datasets for Microglial Motility Analysis Using the MotilA Pipeline.” Zenodo. 10.5281/zenodo.15061566.

Gohlke, Christoph. 2025. “Tifffile: Read and Write Image Data from and to TIFF Files.” Zenodo. 10.5281/zenodo.6795860.

Guizar-Sicairos, Manuel, Samuel T. Thurman, and James R. Fienup. 2008. “Efficient Subpixel Image Registration Algorithms.” Opt. Lett. 33 (2): 156–58. 10.1364/OL.33.000156.

Harris, Charles R., K. Jarrod Millman, Stéfan J. van der Walt, Ralf Gommers, Pauli Virtanen, David Cournapeau, Eric Wieser, et al. 2020. “Array Programming with NumPy.” Nature 585 (7825): 357–62. 10.1038/s41586-020-2649-2.

Helmchen, Fritjof, and Winfried Denk. 2005. “Deep Tissue Two-Photon Microscopy.” Nature Methods 2 (12): 932–40. 10.1038/nmeth818.

Horton, Nicholas G., Ke Wang, Demirhan Kobat, Catharine G. Clark, Frank W. Wise, Chris B. Schaffer, and Chris Xu. 2013. “In Vivo Three-Photon Microscopy of Subcortical Structures Within an Intact Mouse Brain.” Nature Photonics 7 (3): 205–9. 10.1038/nphoton.2012.336.

Kuglin, C. D., and D. C. Hines. 1975. “The Phase Correlation Image Alignment Method.” In Proceedings of the IEEE International Conference on Cybernetics and Society, 163–65. New York, NY, USA.

Lee, Rachel M., Leanna R. Eisenman, Satya Khuon, Jesse S. Aaron, and Teng-Leong Chew. 2024. “Believing Is Seeing – the Deceptive Influence of Bias in Quantitative Microscopy.” Journal of Cell Science 137 (1): jcs261567. 10.1242/jcs.261567.

Li, C. H., and P. K. S. Tam. 1998. “An Iterative Algorithm for Minimum Cross Entropy Thresholding.” Pattern Recognition Letters 18 (8): 771–76. 10.1016/S0167-8655(98)00057-9.

Miles, Alistair jakirkham, Joe Hamman, Dimitri Papadopoulos Orfanos, David Stansby, M Bussonnier, Josh Moore, et al. 2025. “Zarr-Developers/Zarr-Python: V3.0.6.” Zenodo. 10.5281/zenodo.3773449.

Misra, Ishan, C. Lawrence Zitnick, Margaret Mitchell, and Ross B. Girshick. 2015. “Seeing Through the Human Reporting Bias: Visual Classifiers from Noisy Human-Centric Labels.” 2016 IEEE Conference on Computer Vision and Pattern Recognition (CVPR), 2930–39. 10.1109/cvpr.2016.320.

Nebeling, Felix Christopher, Stefanie Poll, Lena Christine Justus, Julia Steffen, Kevin Keppler, Manuel Mittag, and Martin Fuhrmann. 2023. “Microglial Motility Is Modulated by Neuronal Activity and Correlates with Dendritic Spine Plasticity in the Hippocampus of Awake Mice.” Edited by Laura L Colgin and Ania K Majewska. eLife 12 (February): e83176. 10.7554/eLife.83176.

Nimmerjahn, Axel, Frank Kirchhoff, and Fritjof Helmchen. 2005. “Resting Microglial Cells Are Highly Dynamic Surveillants of Brain Parenchyma in Vivo.” Science 308 (5726): 1314–18. 10.1126/science.1110647.

Otsu, Nobuyuki. 1979. “A Threshold Selection Method from Gray-Level Histograms.” IEEE Transactions on Systems, Man, and Cybernetics 9 (1): 62–66. 10.1109/TSMC.1979.4310076.

Pizer, Stephen M., E. Philip Amburn, John D. Austin, Robert Cromartie, Ari Geselowitz, Trey Greer, Bart ter Haar Romeny, John B. Zimmerman, and Karel Zuiderveld. 1987. “Adaptive Histogram Equalization and Its Variations.” Computer Vision, Graphics, and Image Processing 39 (3): 355–68. 10.1016/S0734-189X(87)80186-X.

Prewitt, J. M. S., and M. L. Mendelsohn. 1966. “The Analysis of Cell Images.” Annals of the New York Academy of Sciences 128 (3): 1035–53. 10.1111/j.1749-6632.1965.tb11715.x.

Prinz, Marco, Steffen Jung, and Josef Priller. 2019. “Microglia Biology: One Century of Evolving Concepts.” Cell 179 (2): 292–311. 10.1016/j.cell.2019.08.053.

Ridler, T. W., S. Calvard, and Full name. 1978. “Picture Thresholding Using an Iterative Selection Method.” IEEE Transactions on Systems, Man, and Cybernetics 8 (8): 630–32. 10.1109/TSMC.1978.4310039.

Schindelin, Johannes, Ignacio Arganda-Carreras, Erwin Frise, Verena Kaynig, Mark Longair, Tobias Pietzsch, Stephan Preibisch, et al. 2012. “Fiji: An Open-Source Platform for Biological-Image Analysis.” Nature Methods 9 (7): 676–82. 10.1038/nmeth.2019.

Tremblay, Marie-Ève, Rebecca L. Lowery, and Ania K. Majewska. 2010. “Microglial Interactions with Synapses Are Modulated by Visual Experience.” PLOS Biology 8 (11): 1–16. 10.1371/journal.pbio.1000527.

Virtanen, Pauli, Ralf Gommers, Travis E. Oliphant, Matt Haberland, Tyler Reddy, David Cournapeau, Evgeni Burovski, et al. 2020. “SciPy 1.0: Fundamental Algorithms for Scientific Computing in Python.” Nature Methods 17: 261–72. 10.1038/s41592-019-0686-2.

Wall, Emily, Leslie M. Blaha, Celeste Lyn Paul, Kristin Cook, and Alex Endert. 2018. “Four Perspectives on Human Bias in Visual Analytics.” In Cognitive Biases in Visualizations, edited by Geoffrey Ellis, 29–42. Cham: Springer International Publishing. 10.1007/978-3-319-95831-6_3.

Walt, Stéfan van der, Johannes L. Schönberger, Juan Nunez-Iglesias, François Boulogne, Joshua D. Warner, Neil Yager, Emmanuelle Gouillart, Tony Yu, and the scikit-image contributors. 2014. “Scikit-Image: Image Processing in Python.” PeerJ 2: e453. 10.7717/peerj.453.

Wu, Kensheng, Ekow Otoo, and Arie Shoshani. 2005. “Optimizing Connected Component Labeling Algorithms.” Technical Report LBNL-56864. Lawrence Berkeley National Laboratory, University of California. 10.1117/12.596105.

Yen, Jui-Chang, Fu-Ju Chang, and Shiou-Jyh Chang. 1995. “A New Criterion for Automatic Multilevel Thresholding.” IEEE Transactions on Image Processing 4 (3): 370–78. 10.1109/83.366472.

Zack, Gary W., William E. Rogers, and Samuel A. Latt. 1977. “Automatic Measurement of Sister Chromatid Exchange Frequency.” Journal of Histochemistry & Cytochemistry 25 (7): 741–53. 10.1177/25.7.70454.

